# Stroma secreted IL6 selects for “stem-like” population and alters pancreatic tumor microenvironment by reprogramming metabolic pathways

**DOI:** 10.1101/2020.04.23.041509

**Authors:** Kousik Kesh, Vanessa T Garrido, Austin Doesch, Brittany Durden, Vineet K Gupta, Nikita S Sharma, Michael Lyle, Nagaraj Nagathihalli, Nipun Merchant, Ashok Saluja, Sulagna Banerjee

## Abstract

Pancreatic adenocarcinoma is a devastating disease with an abysmal survival rate of 9%. A robust fibro-inflammatory and desmoplastic stroma, characteristic of pancreatic cancer, contributes to the challenges in developing viable therapeutic strategies in this disease. Apart from constricting blood vessels and preventing efficient drug delivery to the tumor, the stroma also contributes to aggressive biology of the cancer along with its immune-evasive microenvironment. In this study, we show that in pancreatic tumors, the developing stroma increases tumor initiation frequency in pancreatic cancer cells in vivo by enriching for CD133+ aggressive “stem-like” cells. Additionally, the stromal fibroblasts secrete IL6 as the major cytokine, increases glycolytic flux in the pancreatic tumor cells and increases lactate efflux in the microenvironment via activation of the STAT signaling pathway. We also show that the secreted lactate favors activation of M2 macrophages in the tumor microenvironment, which excludes CD8+ T-cells in the tumor. Our data additionally confirms that treatment of pancreatic tumors with anti-IL6 antibody results in tumor regression as well as decreased CD133+ population within the tumor. Furthermore, inhibiting the lactate efflux in the microenvironment reduces M2 macrophages, and makes pancreatic tumors more responsive to anti-PD1 therapy. This suggests that stromal IL6 driven metabolic reprogramming plays a significant role in the development of an immune evasive microenvironment. In conclusion, our study shows that targeting the metabolic pathways affected by stromal IL6 can make pancreatic tumors amenable to checkpoint inhibitor therapy.

## Introduction

Pancreatic cancer has emerged as one of the most aggressive malignancies with a dismal 5-year survival rate of mere 9% ^1^. Aggressive biology, manifested by a high rate of tumor recurrence and resistance to therapy, contributes to poor prognosis in this disease. The pancreatic tumor microenvironment is characterized by the presence of a robust, fibro-inflammatory stroma that is largely immunosuppressive. During tumor progression, this microenvironment develops anatomically distinct “niches”, leading to enrichment of a small population of “stem-like” cells that have a distinct survival advantage over the bulk of the tumor cells. This population contributes to the aggressive biology of the disease ^2, 3, 4^. Our previous studies show that this aggressive population has increased self-renewal properties (contributing to tumor recurrence), increased invasiveness, and ability to evade adverse chemotherapeutic assault on the tumor and is represented by the expression of CD133 on their surface ^5, 6^. Hypoxic regions, nutritional stress and chemotherapeutic agents enrich for this treatment refractory population within the tumor by converting non-aggressive CD133-cells to an aggressive CD133+ population ^7, 8, 9, 10^. This dynamic inter-conversion of non-aggressive population to an aggressive one contributes to a major challenge while developing effective therapy.

The pro-metastatic and pro-tumorigenic role of the pancreatic cancer stroma is well-studied ^11, 12, 13, 14^. In liver and lung cancer, the stromal cells have been reported to reprogram the tumor cells^15^. However, the extent to which the stromal components affect the properties of the tumor is an ongoing area of research. Stroma secreted cytokines like IL6 has been implicated as one of the major inflammatory mediators in pancreatic cancer ^16, 17^. Independent of cytokines, stroma derived metabolites have been shown to promote autophagy in tumor cells and contribute to tumor progression ^18, 19, 20^. While the co-culture of stromal cells with tumor cells is known to enrich for tumor-initiating cells or TIC in other cancers ^21, 22, 23^, whether they promote the enrichment of CD133+ TIC population in pancreatic cancer is not known. Metabolic reprogramming in a tumor cell plays a crucial role for it to develop into an aggressive state that has a high recurrence potential ^24^. Recent research shows that hypoxic microenvironment enriches for CD133+ population within a tumor ^7, 25, 26^. Classically, the hypoxic microenvironment alters cellular metabolism, promoting glycolysis and suppressing mitochondrial respiration. This results in a lower accumulation of ROS in these cells upon treatment with chemotherapy, eventually protecting these cells from apoptosis ^8^. Literature shows that the altered metabolism in the tumor cell has a significant influence on the tumor microenvironment affecting polarization of macrophages to a “pro-tumorigenic” M2 phenotype that in turn can exclude cytotoxic T-cells from entering the tumor, thereby contributing to the immune evasive microenvironment ^27, 28^. However, whether this metabolic reprogramming drives immune suppression in pancreatic cancer is not known. Our recently published studies have shown that tumor intrinsic metabolic pathways like hexosamine biosynthesis pathway can be targeted to modulate the microenvironment and thus sensitize pancreatic cancer cells to checkpoint inhibitor therapy ^29^. Thus, metabolic inhibitors are being evaluated to sensitize normally immune evasive pancreatic tumors to checkpoint inhibitor therapy

In the current study, we show that in pancreatic tumors, the developing stroma increases tumor initiation frequency in pancreatic cancer cells in vivo by enriching for CD133+ aggressive “stem-like” cells. Further, the stromal fibroblasts secrete IL6 as the major cytokine, which in turn, increases glycolysis in the pancreatic tumor cells, leading to increased lactate efflux in the microenvironment. We also show that the secreted lactate as a result of metabolic reprogramming in response to stromal IL6 remodels the tumor immune microenvironment to make it immune evasive, by favoring M2 macrophages in the microenvironment. We show that treatment of pancreatic tumors with anti-IL6 antibody results in tumor regression as well as decreased CD133+ population within the tumor. Similarly, inhibiting the lactate efflux in the microenvironment decreases M2 macrophages and makes notoriously immune-evasive pancreatic tumors more responsive to anti-PD1 therapy.

## Material & Method

### Cell culture, treatment and silencing

MIAPaCa-2 (CRM-CRL-1420) and SU.86.86 (CRL-1837) cell lines were purchased from ATCC and cultured respectively in DMEM (Hyclone) and RPMI 1640 (Hyclone) containing 10% fetal bovine serum (FBS) with 1% Pen Strep (Gibco). Human PSC (ScienCell) cells were maintained in steCM (ScienCell) media according to the recommended conditions. CAFs were isolated from KPC mice according to Dauer et al ^30^ and maintained in DMEM (Hyclone). All the established cell lines were used between passages 5 and 18. All cells were maintained at 37⍰°C in a humidified air atmosphere with 5% CO_2_. 70% confluent cells were used in each experiment. All cell lines were tested for mycoplasma and validated by STR profiling routinely.

Recombinant IL6 (15ug/ml) (Abcam) was used to treat cells in culture for all experiments. For the IL6 receptor silencing, ON-TARGETplus human Il6-R siRNA-SMARTpool (GE Dharmacon) was transfected using HiPerfect transfection reagent according to manufacturer’s protocols. For IL6 receptor, neutralizing experiment, cells were treated with anti-Il6 receptor monoclonal antibody (B-R6, Invitrogen) 1 h before IL6 stimulation. Small molecules inhibitor Stattic and STF31 (Sigma) at indicated concentrations were used in various inhibition experiments. Conditioned media (CM) of tumor cells and PSC was produced using FBS-free basal media to exclude the effects of growth factors in serum for downstream experiments. In normal conditions, 70% confluent cells were cultured in FBS-free basal media for 48 hours. The resulting CM were centrifuged for 10 minutes at 1,000 rpm after collection and stored at −80°C for no more than two months before treating the cells.

### Boyden chamber invasion assay

Boyden chamber invasion inserts (Corning BioCoat) were rehydrated for 2 hours in serum-free medium at 37°C. Cells were plated in the insert, on top of the Matrigel-coated membrane in a serum-free medium. The bottom well contained either PSC-CM or Mia-CM, serving as the attractant. After 24 hours, inserts were washed with PBS, the top of the membrane was scrubbed with a cotton swab to remove any remaining noninvaded cells, fixed in methanol, and stained with crystal violet. Invading cells were counted randomly by microscopy (Magnification, ×20).

### Isolation of CD133-tumor cells

The CD133-population was separated from MIAPaCa-2 cells using MACS separation (Miltenyi Biotech) according to the manufacturer’s protocol. Cells growing in culture were scraped gently into centrifuge tubes and washed once in wash buffer (PBS, 0.5% BSA, 2 mM EDTA), and subsequently incubated with anti-mouse CD133 microbeads for 10 min on ice, and negatively purified for CD133-cells by MACS. The purity of separation was tested for each batch by performing a FACS analysis using Anti-CD133-PE antibody AC141 (Miltenyi Biotech).

### Quantitative real-time PCR and PCR array

Quntatitative real time PCR for SNAIl1, SNAIL2, SNAIL3, ZEB1, TWIST1, HSP70, HSF1, NFKB1, RelA, Suervivin, OCT4, SOX2 and Nanog were performed using primers from Qiagen (QuantiTect primer assay). All data were normalized to 18S (18S QuantiTect Primer Assay; Qiagen).

In another set of experiment, MIAPaCa-2 and SU.86.86 cells were plated and treated in various conditions as described above for quantitative PCR array. The human glucose metabolism array RT2 profiler PCR kit (PAHS-006ZG-4) from QIAGEN was used for RNA expression analysis, according to the manufacturer’s instructions.

### Multiplexed cytometric bead array

Conditioned media from MIA-PACA2, SU86.86 and PSC (Mia-CM, SU-CM and PSC-CM respectively) were collected from 48h cell culture in FBS-free basal media. Murine Pancreatic Stellate cells (PSC) were isolated from normal C57BL6 mice, 1-month old KPC mice ^31^ along with CAFs from fully grown KPC tumors. Isolated cells were grown in tissue culture dishes for 1 week. PSCs were maintained in the inactive form by plating on collagen-coated plates. The conditioned medium from these were collected. The LEGEND plex human inflammation panel 1 (BioLegend 740809) and mouse Th cytokine Panel (BioLegend 740740) were used, and the experiment was performed according to the manufacturer’s instructions.

### Immunofluorescence and immunohistochemistry

Tissues were deparaffinized by heating it at 56°C overnight and then hydrated by dipping it in xylene (15 mins, two times), 100% ethanol,90% ethanol, 70% ethanol (2 times) and PBS 5 mins each. The slides were then steamed with a pH 9 reveal decloaker (BIOCARE Medical) for antigen retrieval, blocked in Dako serum free blocker (Agilent technologies). Primary antibody against αSMA (Abcam), CD133 (Miltenyi), Ki67 (Invitrogen), CD206 (Abcam), CD8 (Abcam) was added overnight. Slides were washed 3X in PBS, secondary antibodies (Alexafluor conjugated) were diluted in SNIPER (BIOCARE Medical) and slides were stained for 30 mins at room temperature. Slides were washed again 3X in PBS and mounted using Prolong Gold anti-fade with DAPI (Thermo Fisher Scientific). Slides were dried overnight and imaged by fluorescent microscope. Images were acquired at 20X and 40X.

### FACS analysis

Single-cell suspensions were prepared from fresh cell culture. Cell fixation and permeabilization was performed with the BD Biosciences Cytofix/Cytoperm kit (BD Biosciences), according to the manufacturer’s instructions. Apoptosis and bromodeoxyuridine (BrdU) incorporation for proliferation was done using Apoptosis and Cell Proliferation Kit following manufacturer’s instructions (BD Biosciences). CD133+ population was analysed by FACS (Miltenyi Biotech, human and mice). All samples were analyzed on BD FACSCANTO II flow cytometers (BD Biosciences). Data was acquired and analyzed with FACS DiVa software (BD Biosciences) and FlowJo Software.

### Sirius red staining and measurements

Tissue sections were stained using picrosirius red staining solution (Chondrex Inc) according to the manufacturer’s instructions. The Sirius red stained area was quantified using ImageJ software by selecting stained fibers in five fields at a magnification of ×100 under a light microscope.

### Western blotting

To analyzed protein expression cells were lysed using RIPA lysis buffer (Boston Bioproducts) containing protease and phosphatase inhibitors (Roche), and protein concentration was estimated using the BCA protein estimation assay (Thermo Scientific). Equal amounts of protein were separated by SDS-PAGE and transferred to nitrocellulose membrane. Blots were probed with antibodies against STAT3 and phospho-STAT3 (Tyr705) (Cell Signaling), after washing re-probed with secondary antibody (Abcam). The bands were visualized using super signal West Dura Extended Duration Subject (34075-Thermo Fisher) in a ChemiDoc (Bio-Rad).

### Activity assay and lactate measurement

Mia-PaCa2and SU.86.86 cells grown in 70% confluent were treated with IL6 in the presence or absence of prior anti IL6-R antibody or static treatment for 24 hrs. Untreated cells were served as a control. Proteins were extracted and equal amount of protein were used for various enzyme activity assay. ALDH, Mitochondria complex4 and PDH enzyme activity assay were performed using the respective assay kits (Abcam) according to manufacturer’s instruction. Total lactate from the cell lysates was measure using Lactate Assay Kit (Sigma, MAK064) according to the manufacturer instruction.

### Extracellular acidification rate (ECAR) determination using Seahorse

MiaPaca-2 cells were plated at a density of 20000 cells in a Seahorse XF96 well plates per well in 80 μL of DMEM media (with 10% FBS) and the plate was kept 1 h at room temperature followed by incubation at 37 °C with 5% CO_2_ for 3 h. Finally, 100 μL of fresh media was added to have total 180 μL per well and incubated for 16 h. Later on, cells were treated with Il6 with or without anti -IL6-R antibody, the culture medium was replaced with glucose-free XF24 Seahorse medium containing 1 mM l-glutamine. Glycolytic flux (basal glycolysis, glycolytic capacity, and glycolytic reserve) were assessed by extracellular acidification rate (ECAR) was measured by the sequential addition of glucose (10 mM), oligomycin (1 μM), and 2-DG (50 mM) in an XF96 Extracellular Flux Analyzer according to the manual of XF glycolysis stress test kit (no. 103020– 100, Seahorse Bioscience). The ECAR values were normalized to the cell counts in each well. Glycolytic capacity was calculated according to the company instruction.

### Glucose uptake assay

Cell-based glucose uptake assay kit (Cayman Chemicals) were used for measuring glucose uptake in Mia-Paca2 and SU86.86 cells. Cells were plated and treated with Il6/ PSC-CM/TNFα in the presence or absence of 30 min prior anti IL6-R antibody/static or STF31 treatment. For labeling, isolated cells were starved in glucose-free media for 30 min prior to labeling with 150 ug/ml 2-NBDG, a fluorescent analog of deoxyglucose. Following incubation for 1 h, the labeled cells were analyzed by flow cytometry (BD Biosciences) or microplate reader (Spectramax-ix3) according to manufacturer’s instruction.

### Animal experiment

Male athymic nude mice and C57Bl6 mice between the ages of 4-6 weeks were purchased from the Jackson Laboratory, Bar Harbour, ME, USA. For tumor initiation experiment, the subcutaneous mice tumor model was exploited. CD133 □ MIA-Paca-2 cells were implanted with human PSC cells in a ratio of (1:9) in the right flank of the mice. Cells were implanted at a limiting dilution manner such as (100:900, 1000:9000 and 10000:90000). Corning® Matrigel® Growth Factor Reduced (GFR) Basement Membrane Matrix, purchased from Corning, Inc, Corning NY, USA and 1X PBS at a ratio of 1:1 were used as a suspension medium for the cells. The appearance of detectable tumor nodule were observed 3 times in a week. The days of tumor appearance were noted in each group. Tumor initiation frequency was calculated using ELDA assay.

To see the effect of PSC in TIC plasticity, CD133-MIA-Paca-2 cells were implanted with human PSC cells in a ration of (1:9) in male athymic nude mice. Tumor progression were monitor by measuring the tumor volume weekly. All mice were sacrificed at the end of the experiment and the tumor was subjected to flow cytometry analysis for CD133 enrichment.

Both the transgenic mouse model of spontaneous PDAC and the orthotopic PDAC mouse model were also used in our study. KPC animals were generated by crossing Lox-Stop-Lox (LSL) KRas^G12D^; Trp53^R172H^ animals with Pdx-1 Cre animals. For IL6 receptor neutralizing experiment, the orthotopic PDAC mouse model was used. 2,000 KPC tumor cells with 18000 CAF cells were suspended in Matrigel (BD Biosciences) and injected in the pancreas of 16 C57BL/6 mice. After 10 days mice were randomly divided in to 2 groups: Group1 received 500 μg of anti-IL-6RAb (Bio-X-cell) in every third day for 4 weeks. While mice received isotype control Ab serve as a control. All the mice were sacrificed and tumor volume and weight were measured. In *In vivo*, STAT3 inhibition was assessed using *Ptf1a*^*Cre*^, *LSL-Kras*^*G12D/+*^, *Tgfbr2*^*flox/flox*^ (PKT) transgenic mice, which rapidly develop pancreatic tumors at 4 weeks of age^32^. In brief, PKT mice were treated with vehicle or the STAT3 inhibitor Ruxolitinib (12 mg/kg/day, three times weekly) by oral gavage for 2 weeks prior to sacrifice and histologic tissues obtained for further analysis.

For inhibition of carbonic anhydrase with WBI-5111 (aka SLC-0111)^33^, KPC and CAF cells were implanted orthotopically in the pancreas of C57BL6 mice in a ratio of 1:9. Treatment with 100mg/kg CA9 inhibitor, WBI-5111 (a gift from Welichem Biotech Inc.) was started 7 days following implantation. 3 injections of 100ug anti-PD1 antibody was administered at Day 17, 18 and 21. Animals were sacrificed 30 days after start of treatment with CA9 inhibitors.

### Statistical analysis

Data were presented as the means ± SEM. Statistical analyses were performed using GraphPad Prism, version 7.0. Differences between two groups were analyzed by Student’s t test. *P* < 0.05 was considered statistically significant. The experiments were repeated at least in triplicate for 3 times.

### Ethics statement

All animal studies were performed according to the protocols approved by IACUC at the University of Miami, USA, in accordance with the principles of the Declaration of Helsinki. All authors had access to all data and have reviewed and approved the final manuscript.

## Results

### Stroma promotes tumor “initiation” and enrichment of self-renewal gene expression

Tumor microenvironmental niches have been implicated in enriching for “cancer stem cells” or “tumor-initiating cells” in a number of cancers. Previously published studies have revealed that hypoxia enriches for pancreatic tumor-initiating cells that are aggressive and highly metastatic ^7^. To study if the stromal fibroblasts affected an enrichment of CD133+ population, we treated human pancreatic cancer cells MIA-PaCa2 and SU86.86 with conditioned media from human pancreatic stellate cells (hPSC). Our results showed a distinct enrichment of CD133+ population in both cells lines (Figure 1A, B). Treatment with PSC conditioned media also increased the invasiveness of CD133^lo^ cells *in vitro* (as seen by Boyden chamber assay) (Figure 1C) as well as increased expression of genes involved in cancer metastasis (Figure 1D). Additionally, treatment with PSC conditioned media also upregulated pro-survival genes like HSP70, HSF1, NF-kB and Survivin (Figure 1E), indicating an enrichment of aggressive, metastatic cells with enhanced survival advantage. To further see if stroma promoted tumorigenesis *in vivo*, we implanted CD133-MIA-PaCa2 cells with and without PSC in a 1:9 ratio. Our results showed that while all animals in the set that was implanted with the PSCs had tumor initiation by day 37, the set of animals lacking PSC 3/5 animals failed to form tumors by the end of study at Day70 (Figure 1F). Furthermore, we observed an increased CD133+ population in the tumors implanted with the PSCs compared the those that lacked PSC (Figure 1G). Calculation of tumor initiation frequency by ELDA ^34^ showed an almost 6-fold increase in the presence of stroma (Figure 1H). Additionally, tumors lacking PSC showed a delayed tumor growth compared to the ones that had PSC co-implanted with them (Figure 1I).

**Figure 1.**
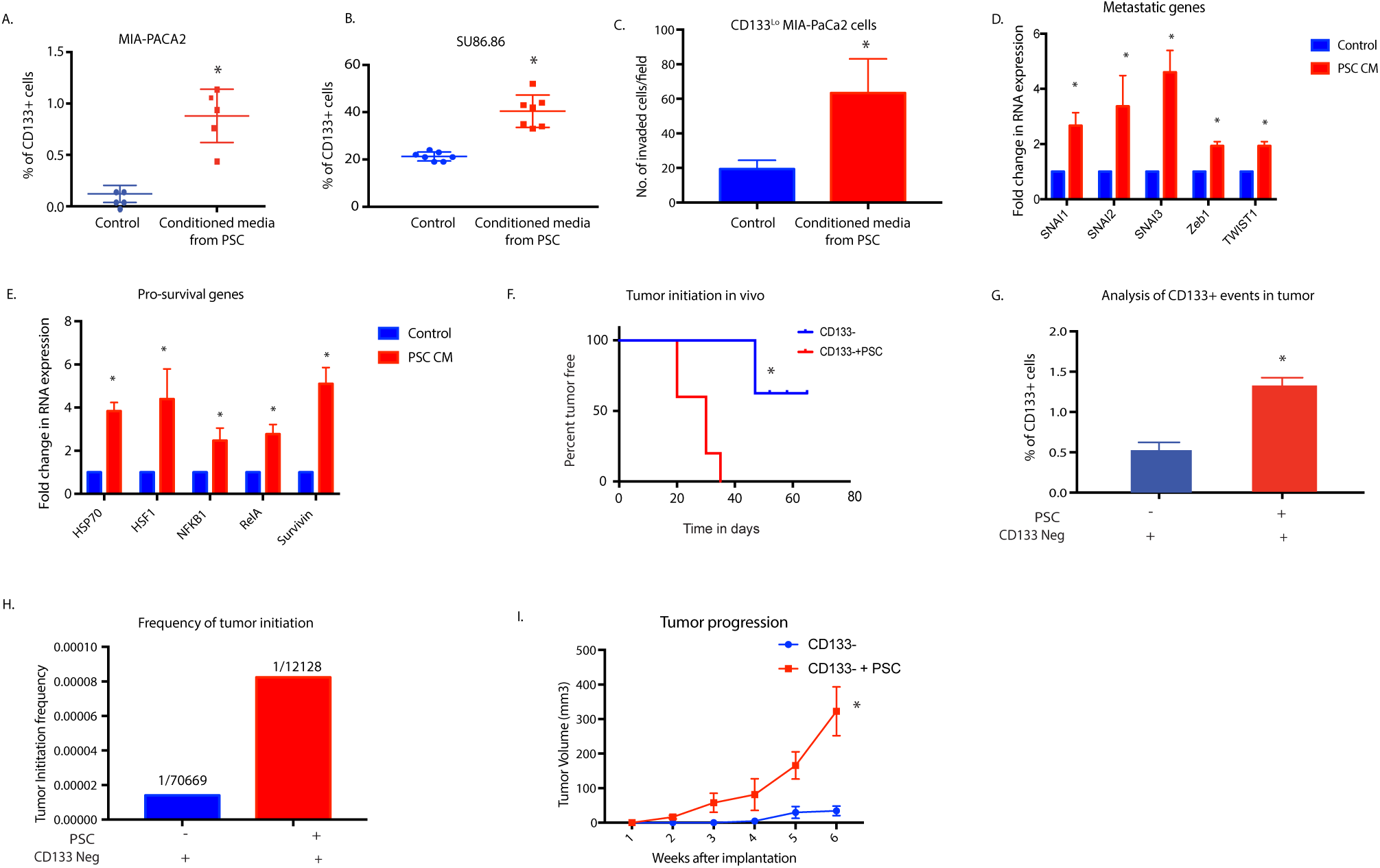
Stroma enriches for CD133+ population in pancreatic cancer: Treatment of human pancreatic cancer cells MIA-PACA2 (A) and Su86.86 (B) with human pancreatic stellate cell (hPSC) conditioned media showed an increase in the CD133+ population. PSC conditioned media also increased invasive potential of CD133^Lo^ pancreatic cancer cell MIA-PACA2 cells (C) along with increasing expression of genes involved in metastasis (D). Additionally, treatment with PSC conditioned media also increased expression of pro-survival genes in MIA-PACA2 cells €. Presence of stroma increased tumor initiation potential in CD133-cells (F) along with increasing % of CD133+ cells in the tumor (G). Extreme Limiting Dilution Assay to calculate the frequency of tumor initiation showed a 6-fold increase in tumor initiation potential in the presence of stroma (H). Presence of stroma also increased tumor progression rate (I). * represents significant (p<0.05) changes compared to untreated.

### Enrichment of CD133+ aggressive population in the presence of stroma is mediated by stromal IL6

Since treatment with PSC conditioned media enriched for an aggressive population and co-implanting CD133^lo^ cells with PSC led to increased tumor initiation, we next analyzed the PSC conditioned medium for secreted factors. Our multiplexed analysis showed that the PSC conditioned media had a significant high IL6 and TNFα in it (Figure 2A). To study if the pancreatic stroma produced these cytokines during tumor progression, we next isolated PSC from the pancreas of the Kras^G12D^; TP53^R172H^; Pdx-Cre or KPC mouse model of pancreatic cancer as a pre-cancerous stage (i.e from 1 month old animals), from the mid-cancer stage (i.e when the PSCs start getting activated) and from full tumor-bearing animals (when the PSC transform to Cancer-Associated Fibroblasts). An analysis of the secretion of cytokines from these cells showed a similar increase in IL-6 and TNFα as the tumor progressed (Figure 2B). The histology of the KPC animal-derived tumors showed progressive development of stroma (observed by αSMA staining) at these stages (Figure 2C). Since the predominant cytokine produced by the stromal fibroblasts was IL6 and TNFα, we next treated the MIA-PACA2 and SU-86.86 cells with IL6 and TNFα. While treatment with IL6 showed a significant increase in CD133 population (Figure 2D, E), treatment with TNFα did not show any significant enrichment (Supplementary Figure 1A). We thus focused our effort on IL6. To test the specificity of IL6 in enriching for CD133+ population, we treated the cells with Anti-IL6 Ab for 1h before treating with IL6. Our results showed that IL6 mediated increase in CD133+ population was decreased in the presence of Anti-IL6 Ab, indicating that this was enrichment was being orchestrated by IL6. Similarly, when these cells were pretreated with Anti-IL6 Ab and then exposed to PSC conditioned media, there was a significant decrease in the CD133+ population (Figure 2D, E). Additionally, IL6 mediated increase in expression of self-renewal genes was decreased when cells were pre-treated with Anti-IL6 Ab (Figure 2F). Aldehyde dehydrogenase activity is associated with an aggressive and “stem-like” population represented that are CD133+ ^35^. Our results showed an increase in Aldh activity that was decreased upon treatment with anti-IL6 Ab (Figure 2G). Consistent with this, pre-treatment of pancreatic cancer cells with IL6 also increased their resistance to gemcitabine and reduced apoptosis in these cells (Supplementary Figure 2).

**Figure 2:**
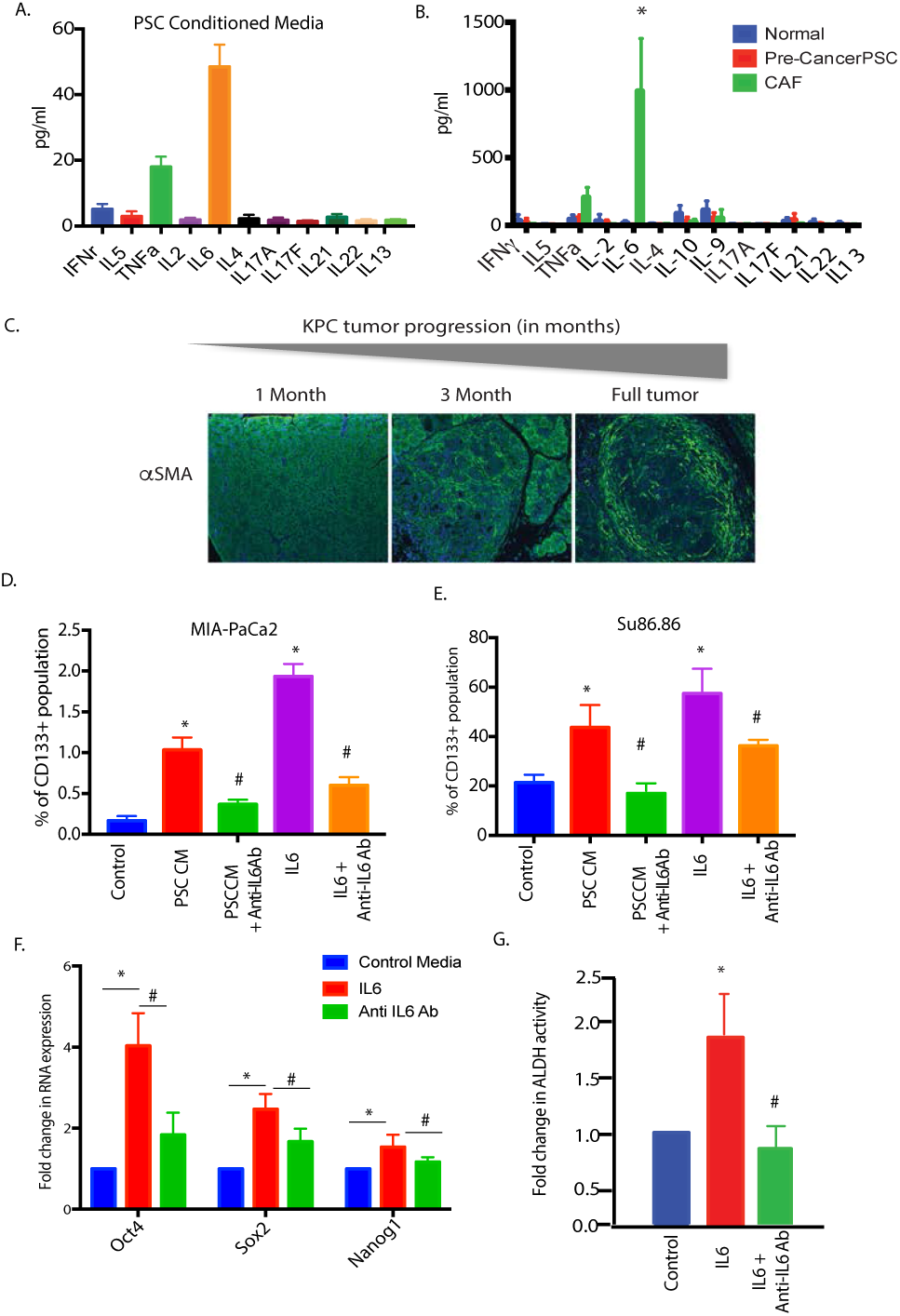
IL6, the primary cytokine produced by stromal cells enriches for CD133+ population: Analysis of PSC conditioned media showed IL6 to be the main cytokine produced by the PSC cells in culture (A). Similar secretion of IL6 was observed when normal, pre-cancer PSC and CAFs from full tumors were analyzed for secreted cytokines (B). Pancreatic cancer stroma develops as the tumor progresses as observed by the increased expression of alpha SMA in tumors (C). Increase in CD133+ cells upon IL6 treatment or PSC treatment was observed to be reduced with IL6 signaling was blocked using anti-IL6R antibody in both MIA-PACA2 (D) and SU86.86 cells (E). Treatment with IL6 increased stemness genes (F) and ALDH activity (G), which were reversed when signaling was blocked with anti-IL6R antibody. * represents significant (p<0.05) changes compared to untreated and # represents significant (p<0.05) changes compared to IL6 treatment.

### Stromal IL6 increases glucose uptake and flux through glycolysis in enriched CD133+ cells

Previously published results from our laboratory showed that CD133+ cells had increased metabolic flux through the glycolytic pathway and a very rudimentary oxidative phosphorylation ^8^. This contributed to the upregulation of anti-apoptotic and pro-survival genes in this population. To determine if the IL6 secreted by CAFs altered the metabolic profile of the cells as they enriched for CD133+ cells, we performed an energy phenotyping assay on SU86.86 and MIA-PACA2 cells using a Seahorse platform. Our results showed that upon treatment with IL6, both cell lines had increased extracellular acidification rate or ECAR (Figure 3A, B). Further, glycolytic capacity (Figure 3C) was increased with IL6 in both cell lines and decreased when signaling was blocked by with anti-IL6 antibody. We next confirmed this by estimating the lactate secreted by the cells. Our results showed that increased lactate was produced by tumor cells in the presence of IL6. This was decreased when IL6 was inhibited using anti-IL6 antibody (Figure 3D). We next determined if the gene expression of the glucose utilization enzymes were affected by stromal IL6 by using a PCR array. Our results showed that treatment with stromal IL6 increased genes involved in glucose metabolism, specifically glycolysis, that decreased when IL6 receptor on tumor cells was silenced by siRNA for IL6R in SU86.86 (Figure 3E,F). Silencing of IL6R was confirmed by qRTPCR (Supplementary Figure 1B). To determine if increased glycolysis and thus ECAR in the presence of stromal IL6 is mediated by increased glucose transporter activity, we measured glucose uptake (using fluorescent glucose analog 2-NBDG) by the tumor cells after blocking IL6 receptor (using anti IL6R antibody) and glucose transporter inhibitor STF31. Our results showed increased glucose uptake in the presence of both PSC-CM and IL6 treatment in Su86.86 cells, which was abrogated by treatment with anti-IL6 antibody (Figure 3 G, H). Similar results were observed in MIA-PaCa2 cells as well (Supplementary Figure 1C). We further observed that the effect of IL6 was reverted upon silencing the IL-6 receptor on tumor cells (by siRNA for IL6R) as well as upon inhibition of glucose transporters by STF31 (Figure 3I). This indicated that stromal IL6 mediated increased glycolysis and ECAR in pancreatic tumor cells by increased glucose uptake via glucose transporters.

**Figure 3.**
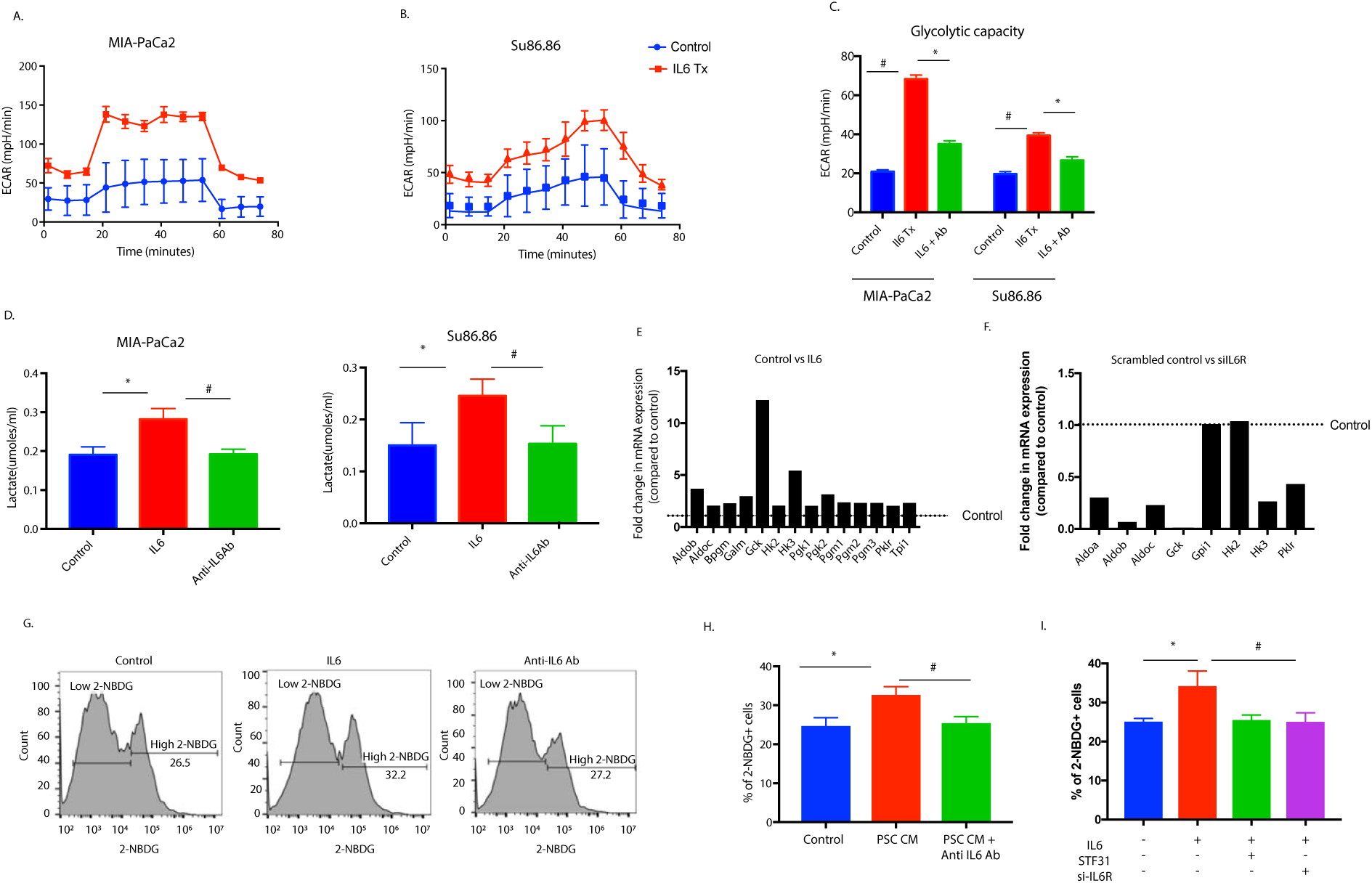
Treatment with IL6 alters metabolic profile in pancreatic cancer cells: Treatment with IL6 increased ECAR in MIA-PACA2 (A) and Su86.86 cells (B) and increased the glycolytic capacity of the cells (C). An increase in lactate production was also observed in both MIA-PACA2 and SU86.86 cells (D), that was reduced when IL6 signaling was blocked. Expression of genes involved in glucose metabolism was altered upon IL6 treatment (E). Similarly, blocking IL6 signaling by silencing IL6 receptor showed a downregulation of glucose metabolism genes (F). IL6 also increased glucose uptake as observed by 2-NBDG uptake assay by flow cytometry (G). Glucose uptake was similarly increased with SU86.86 cells were treated with PSC conditioned media (H). This effect was reverted upon blocking IL6 signaling. Additionally, inhibition of glucose transporter using 5uM STF31 or by silencing IL6R decreased the % of 2-NBDG+ cells. * represents significant (p<0.05) changes compared to untreated and # represents significant (p<0.05) changes compared to IL6 treatment.

### Stromal IL6 induces metabolic reprogramming and enrichment of CD133+ population via STAT3 signaling

To study if stromal IL6 was activating the canonical JAK/STAT pathway in the tumor cells, we next evaluated the activation of STAT3 by treating with IL6. Our studies showed STAT3 was activated (as seen by its phosphorylation as early as 1h and remained activated for 6h (Figure 4A). Similar phosphorylation of STAT3 was observed with PSC conditioned media as well (Figure 4B, Supplementary Figure 3A). We next evaluated if the enrichment of CD133+ population as well as the metabolic changes observed in pancreatic cancer cells, were dependent on activation of STAT3. To determine this, we stimulated pancreatic cancer cells with IL6 in the presence of a STAT3 inhibitor Stattic. Our results showed that inhibition of STAT3 signaling in the presence of IL6 decreased CD133+ population (Figure 4C, Supplementary Figure 3B), and Aldh activity (Figure 4D). Additionally, treatment with 1μM Stattic in the presence of IL6 decreased glucose uptake (Figure 4E, Supplementary Figure 3B) as well as lactate production (Figure 4F, Supplementary Figure 3C) in pancreatic cancer cells Su86.86 and MIA-PACA2. In vivo, treatment with Ruxolitinib, an inhibitor of STAT signaling pathway also reduced CD133+ cells in pancreatic cancer tissues from PKT mice (Figure 4G).

**Figure 4.**
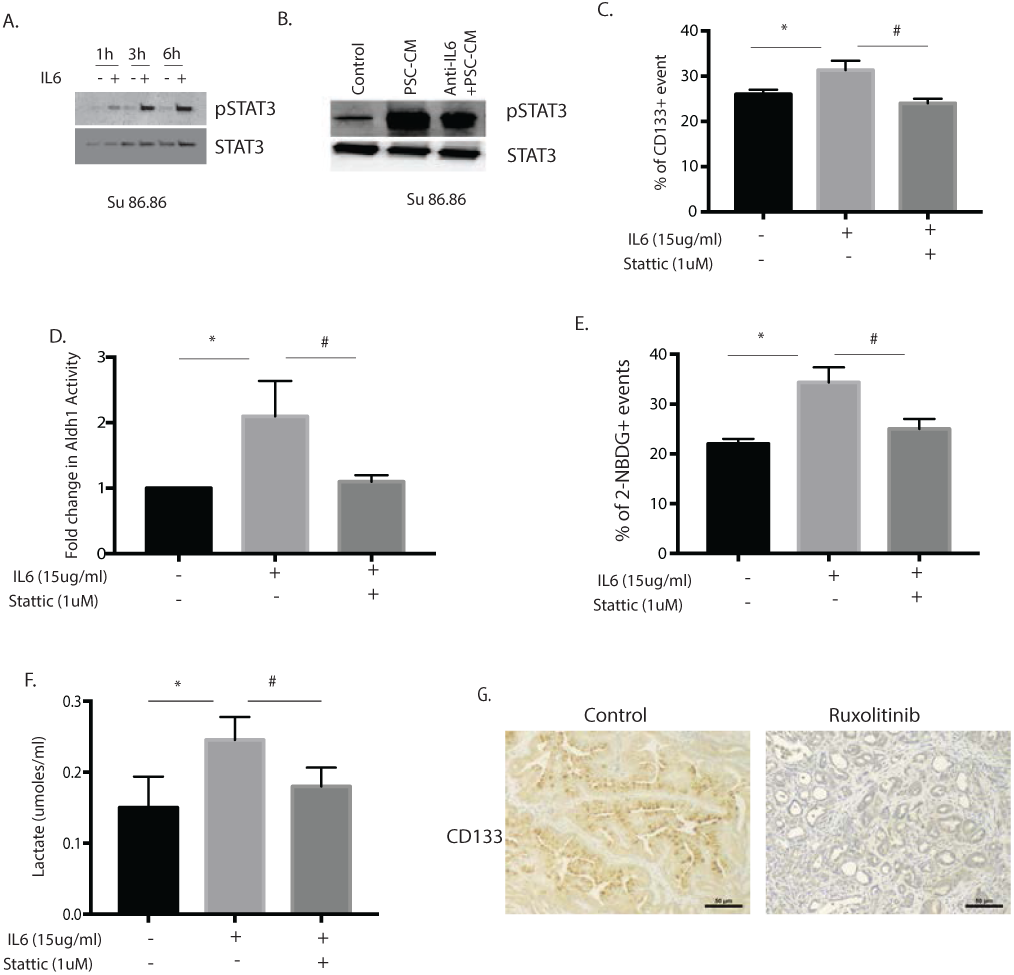
IL6 induced increase in stemness and metabolic changes are mediated via STAT3 signaling. Treatment with IL6 (A) or PSC conditioned media (PSC-CM) induced activation of STAT3 in SU86.86 cells (B). IL6 induced increase in CD133+ population (C) ALdh1 activity (D), glucose uptake (E) and extracellular lactate (F) was reduced when STAT3 signaling was blocked by small molecule inhibitor Stattic. Treatment of pancreatic tumors in PKT mice with Ruxolitinib, a STAT signaling inhibitor also decreased CD133+ cells in the tumor tissues as seen in the representative pictures (G). * represents significant (p<0.05) changes compared to untreated and # represents significant (p<0.05) changes compared to IL6 treatment.

### Stromal IL6 prevents glucose from entering oxidative phosphorylation

Our gene expression analysis following IL6 treatment and an IPA analysis revealed that the increased glycolysis was also associated with a decreased transcription of some of the mitochondrial activity genes like PDHA, KGDH, aconitase and SDHA-D (Figure 5A). One of the proteins that regulate the entry of glycolytic intermediates to mitochondria and oxidative phosphorylation is PDH1. To determine if treatment with IL6 decreased the activity of PDH1, this further, we estimated the activity of PDH1 SU86.86 cells. Our result showed that while treatment with IL6 decreased PDH1 activity, pre-treatment with anti-IL6R or Stattic reverted this inhibition (Figure 5B,C). In MIA-PaCa2 cells; however, this effect was not as pronounced as in Su86.86 cells (Supplementary Fig 3D), IL6 decreased PDH1 activity, however, pretreatment with anti-IL6R and/or static showed a modest recovery. In addition, IL6 treatment inhibited mitochondrial complex 1 activity, which was recovered once signaling was blocked by IL6Ab in both MIA-PaCa2 and Su.86.86 cells (Figure 5D, E). This indicated that stromal IL6 increased glycolysis in cancer cells and suppressed the entry of glucose to oxidative phosphorylation.

**Figure 5.**
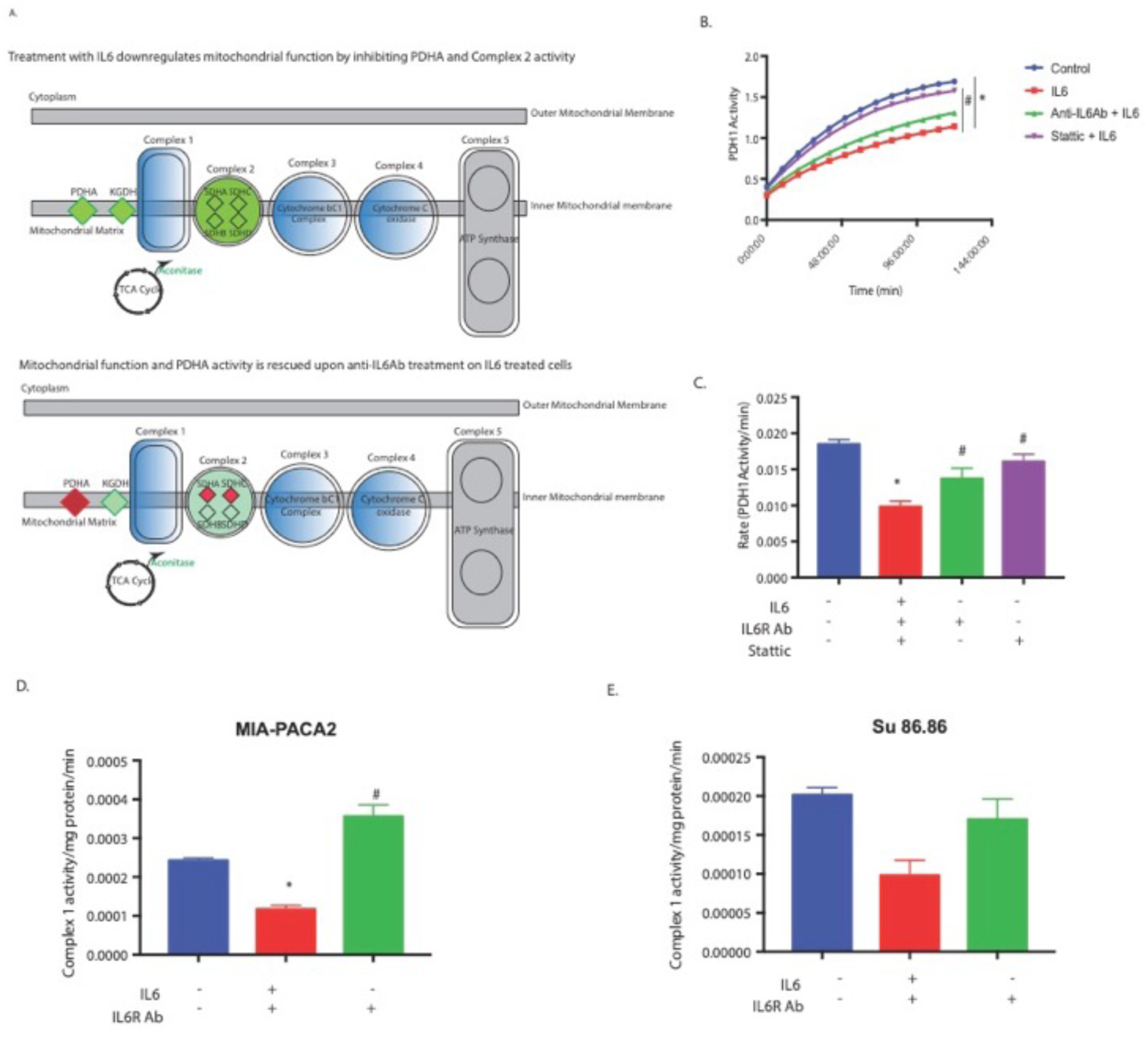
Stromal IL6 prevented entry of glucose into mitochondria: Gene expression analysis showed decrease in mitochondrial “gatekeeper” PDHA and Mitochondrial Complex II genes (A). Further, treatment with IL6 showed decreased PDH1 (PDHA) activity which was rescued when IL6 signaling or STAT3 signaling was blocked by anti IL6R antibody and Stattic respectively (B, C). Complex 1 activity was also decreased with IL6 in both MIA-PACA2 (D) and Su86.86 cells (E), that was rescued upon blocking IL6 signaling with anti IL6R antibody. * represents significant (p<0.05) changes compared to untreated and # represents significant (p<0.05) changes compared to IL6 treatment.

### Inhibition of IL6 inhibits pancreatic tumor progression

IL6 has been shown to be involved in progression of pancreatic tumors by others ^16, 36, 37, 38, 39^. Furthermore, in pancreatic cancer patient population, increased expression of IL6 correlated with poor survival (Figure 6A). To confirm this in our animal model, we next implanted KPC001 cells with CAF cells orthotopically in the pancreas of the C57Bl6 mice in a ratio of 1:9. 7 days following implantation, the animals were treated with anti-IL6 Ab. After 30 days of treatment, our results showed a significant decrease in tumor volume (Figure 6B) and tumor weight (Figure 6C). Further, the analysis of the histology of these slides further showed a decrease in CD133+ population in the Anti-IL6 treated group (Figure 6D). In addition, H&E and Ki67 staining showed a general decrease in proliferative cells (Figure 6E). We next evaluated the collagen deposition in the pancreatic tumor microenvironment. Treatment with anti-IL6 antibody showed significant depletion of collagen in the tumors (Figure 7A) compared to the control animals (Figure 6F).

**Figure 6:**
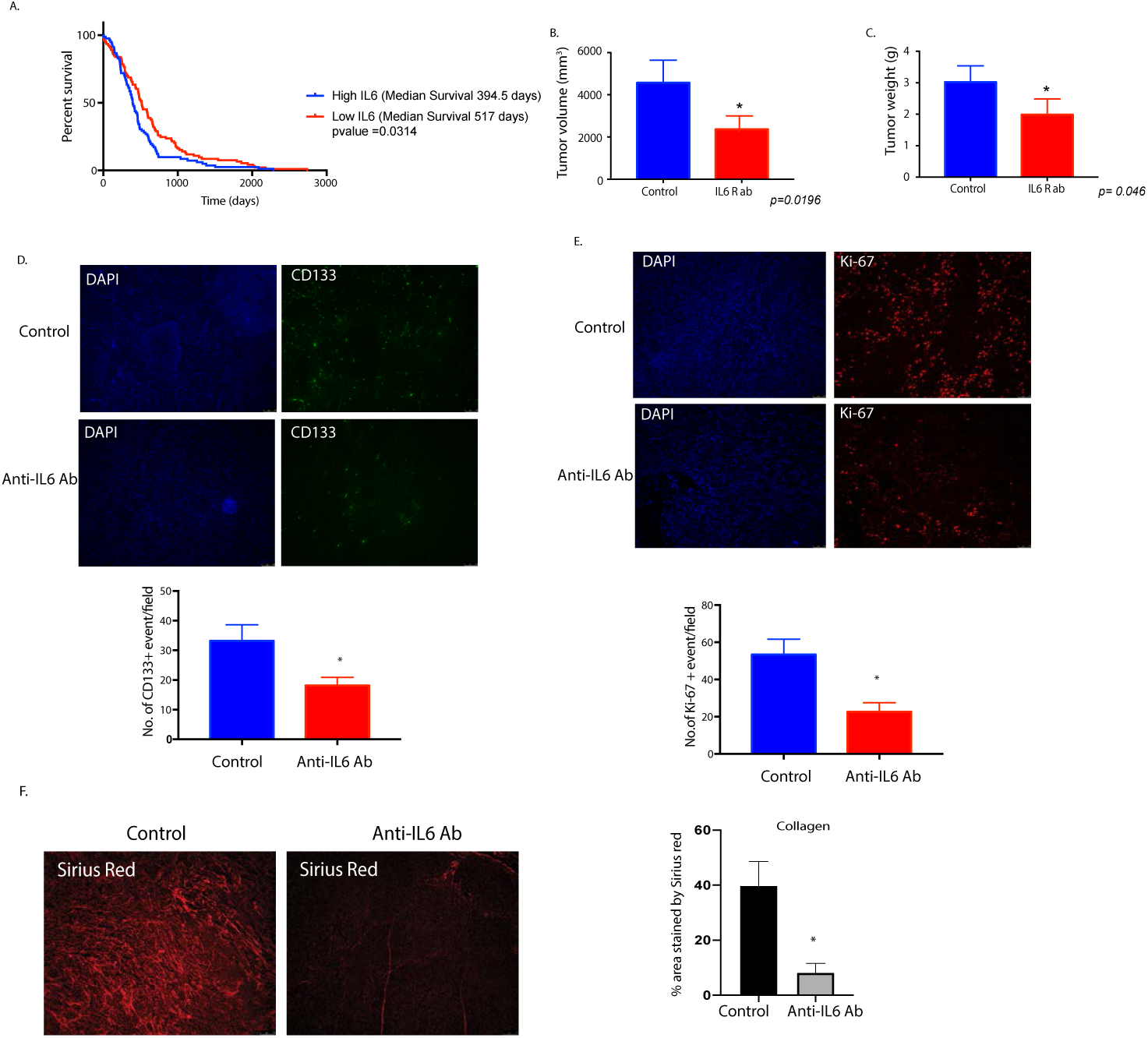
Treatment with IL6 neutralizing antibody causes pancreatic tumor regression. TCGA database shows IL6 expression in pancreatic cancer patients correlate with poor survival (A). Treatment of orthotopically implanted pancreatic cancer cells with IL6 neutralizing antibody showed decrease in tumor volume (B) and tumor weight (C). Immunofluorescence showed decrease in CD133 (D) and Ki67 (E) events. Treatment with IL6 also decreased collagen deposition in the tumors as seen by picrosirius red stain (F). * represents significant (p<0.05) changes compared to untreated samples.

**Figure 7.**
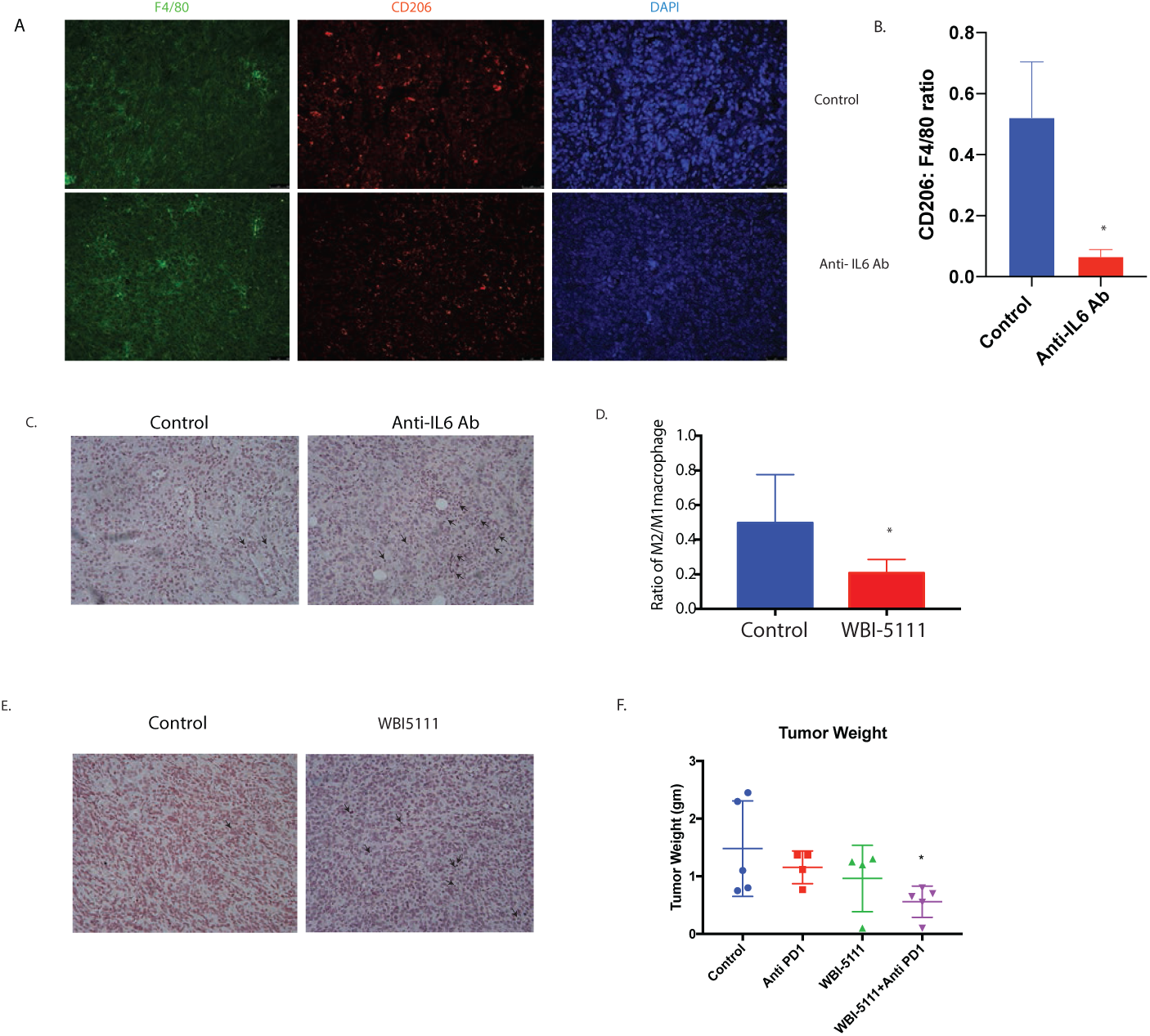
Inhibition of IL6 and effluxed lactate promote CD8+ infiltration and sensitize pancreatic tumors to anti-PD1 therapy. Treatment with IL6 neutralizing antibody decreased M2 activated macrophages ratio in the pancreatic tumor microenvironment (A, B) and increased CD8+ infiltration (indicated by black arrow) (C) in the tumor microenvironment. Inhibition of efflux of lactate using a small molecule inhibitor WBI-5111 that inhibits CA9 showed similar decrease in M2:M1 macrophage ratio in vivo (D). Treatment with WBI-5111 also increased CD8+ infiltration in the microenvironment as indicated by black arrow (E) and sensitized pancreatic tumors to anti-PD1 therapy.

### Inhibition of the IL6 as well as effluxed lactate by CA9 inhibitor modulated pancreatic tumor microenvironment

To determine the effect of IL6 inhibition on macrophage polarization, we next stained the anti IL6 Ab treated tumors with pan macrophage marker F4/80 and M2 marker CD206. A ratio of CD206:F4/80 revealed that anti-IL6 treated tumors had decreased M2 macrophages compared to the control animals (Figure 7A). Along with these, there was an increased infiltration of CD8+ T-cells in the tumors when IL6 signaling was blocked (Figure 7B, Supplementary Figure 4A).

To further study if it was the effluxed lactate in the pancreatic tumor that contributed to the increased CD8+ infiltration, we next implanted KPC:CAF cells in the pancreas of C57BL6 animals and treated them with a CA9 inhibitor. CA9 promotes efflux of lactate from cells to prevent a lactate build up. Analysis of tumors by flow cytometry showed an increased M1/M2 ratio of macrophages within the tumor (Figure 7D). Additionally, the analysis of CD8+ population within the tumors indicated an increase in the CD8+ T-cells (Figure 7E, Supplementary Figure 4B). IL6 inhibition has been shown to sensitize pancreatic tumors to immune checkpoint inhibitor therapy ^40^. To study if inhibition of lactate efflux affected the response of pancreatic tumors to anti-PD1 therapy, we next evaluated the effect of CA9 inhibitor in combination with anti-PD1 therapy. Our results showed that this combination was significantly better in decreasing pancreatic tumors than either therapy alone (Figure 6F).

## Discussion

Many studies have defined the “stem-like” CD133+ population as the “Tumor Initiating Cell or TIC population. However, how this population arises in a tumor has remained enigmatic. Our studies show that CD133+ population in a pancreatic tumor can form tumors at very low dilutions ^5^. Further, overexpression of CD133 in a cell line that lacks the expression of this gene (MIA-PACA2) results in the induction of self-renewal properties in these cells ^6^. This establishes that CD133+ population is an accurate representation of the aggressive treatment-refractory population in pancreatic cancer. By definition, TICs have the ability to form tumors that recapitulate the heterogeneity of the primary tumor from which they were isolated following orthotopic transplantation into mice ^41, 42^. As the progeny of TICs differentiate, they lose their ability to initiate tumors, despite their identical genetic landscape and become “non-TIC” population. However, studies indicate these CD133+ populations are not necessarily the “cell of origin” for pancreatic cancer and thus tumor-initiating cell *per se* but rather a population that is responsible for recurrence, owing to an increased survival advantage. This population of tumor cells has increased adaptation to unfavorable microenvironment like hypoxia and nutrient deprivation. These heightened adaptations to stress in this CD133+ population gives them a distinct advantage over the CD133-population in tiding through unfavorable conditions until the host-environment is conducive for tumor relapse at the primary or metastatic site. Thus, in essence, these cells are the “roots” of aggressive tumors for which there is currently no effective treatments ^4^. Our current study shows that in the presence of stroma, there is an enrichment of CD133+ population within pancreatic tumors (Figure 1).

The role of the stromal cells in the pancreatic tumor microenvironment has gained importance in the last decade. Yet, how the stroma influences the properties of the tumor has remained unknown. A study by Ohlund et al showed the heterogeneity in the cancer-associated fibroblasts (CAFs) in pancreatic tumors, classifying them into inflammatory CAFs (iCAFs) and myofibroblastic CAFs or myCAFs ^16^. This study also showed that inflammatory CAFs secreted IL6 as one of the major cytokines. This is consistent with our findings that during the progression of pancreatic cancer, as the stromal develops and possibly differentiates into iCAFs and myCAFs, there is increased secretion of IL6 (Figure 2). Furthermore, our results show that the stroma secreted IL6 activates the STAT signaling pathway to enrich for “stem-like” CD133+ cells as well as alter the metabolic profile of the pancreatic cancer cells (Figure 3,4). When IL6 signaling was inhibited by blocking the IL6 receptors (by either anti-IL6R antibody or by si-IL6R), a reversal of the enrichment as well as metabolic reprogramming was observed. A similar reversal was also observed when STAT3 was inhibited by inhibitor Stattic (Figure 4). Interestingly, an analysis of STAT3 target genes (in the CHEA transcription factor target database) revealed that stemness genes like Sox2, Nanog and Oct4 as well as certain metabolic genes like Carbonic anhydrases and LDH were targets of STAT3 transcription factors. This indicated that stromal IL6, activated STAT signaling pathway to upregulate mRNA expression of both stemness genes as well as metabolic genes in pancreatic tumor cells (Supplementary Figure 4C).

IL6 induced metabolic reprogramming in pancreatic tumor cells. We observed increased glycolysis and reduced mitochondrial activity in cells treated with IL6. Interestingly, IL6 resulted in increased production of lactate and inhibited the activity of pyruvate dehydrogenase or PDH1, a gatekeeper enzyme of mitochondrial respiration. This was further confirmed upon assaying for mitochondrial complex 1 activity. Treatment with IL6 decreased mitochondrial complex 1 activity confirming that the pancreatic tumor cells were preferentially synthesizing lactate in the presence of IL6. Lactate is effluxed in the microenvironment by activity of CA9.

Studies have shown that this acidification of the microenvironment contributes to immune suppression by favoring polarization of macrophages to an M2 phenotype, which in turn excludes CD8+ T-cells ^27^. It is well known that pancreatic tumors have an immune suppressive microenvironment with limited CD8+ T-cells. However, whether this is because of effluxed lactate is not known. Our study showed that inhibiting IL6 in vivo using an IL6R antibody or by inhibiting the efflux of lactate by inhibiting CA9, decreased the M2 macrophages in the microenvironment, and increased CD8+ infiltration (Figure 7). Remarkably, inhibiting IL6 had a striking effect on collagen in the stroma (Figure 7A). A recently published study from our laboratory show that inhibiting of hexosamine biosynthesis pathway decreased collagen in the pancreatic tumors and increased CD8+ infiltration ^29^. Whether inhibition of IL6 signaling functions similarly is unclear and beyond the scope of the current manuscript. Increased infiltration of CD8+ cells can potentially sensitize tumors to immune therapy. Further, previous research has shown that inhibition of IL6 can sensitize pancreatic tumors to immune therapy ^40^. To study if this overcoming of immune evasion by inhibition of IL6 signaling is due to decreased lactate efflux by tumor cells, we inhibited lactate efflux by inhibiting CA9. Our results showed that this sensitized pancreatic tumors to anti-PD1 therapy (Figure 7). This is a significant observation in the context of pancreatic cancer since this disease is notoriously resistant to immune therapy.

Overall, the current study shows the importance of stromal cells in promoting tumor progression by enriching for an aggressive and stem-like population within the tumor. It further shows that stroma can drive a metabolic reprogramming and favor glycolysis in the tumor cells to promote increased efflux of lactate in the microenvironment, that can contribute to the immune evasive property of the tumor. Finally, our study shows that we can target the IL6 induced lactate efflux from tumor to revert the immune evasive microenvironment to an immune-supportive one and thus augment outcomes of checkpoint inhibitor therapy.

