## Supplementary figures for "Stroma secreted IL6 selects for “stem-like” population and alters pancreatic tumor microenvironment by reprogramming metabolic pathways"

Supplementary Figure Legends

Supplementary Figure 1: Treatment with TNF-alpha did not alter glucose uptake (as measured by 2-NBDG staining) in MIA-PACA2 or SU8686 staining (A). Validation of IL6R silencing in pancreatic cancer cell Su86.86 (B). 2NBDG assay  in MIA-PACA2 cells (C).

Supplementary Figure 2: Treatment of pancreatic cancer cells (BxPC3 and CFPAC ) with IL6 increased their resistance to Gemcitabine induced apoptosis.

Supplementary Figure 3: Activation of STAT3 following treatment with PSC conditioned media on MIA-PACA2 (A). Inhibition of STAT3 signaling by Stattic decreased IL6 induced CD133+ population and lactate production. PDH1 activity assay (kinetics and rate of activity) in MIA-PACA2 cells following IL6 treatment, blocking IL6 signaling by anti IL6R antibody and static respectively.

Supplementary Figure 4. Quantitation of infiltrated CD8+ T cells in IL6 neutralizing antibody treated tumors (A) and WBI-5111 treated tumor (B). Target genes of STAT3 as observed in the CHEA transcription factor target database (C)

### Supplementary Figures

A.

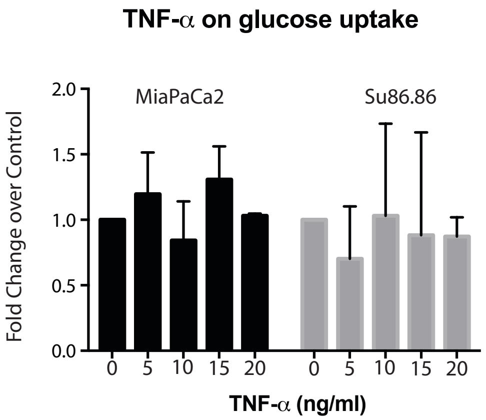
TNF-α on glucose uptake
Fold Change over Control
MiaPaCa2
Su86.86
2.0
1.5
1.0
0.5
0.0
0 5 10 15 20 0 5 10 15 20
TNF-α (ng/ml)

B.

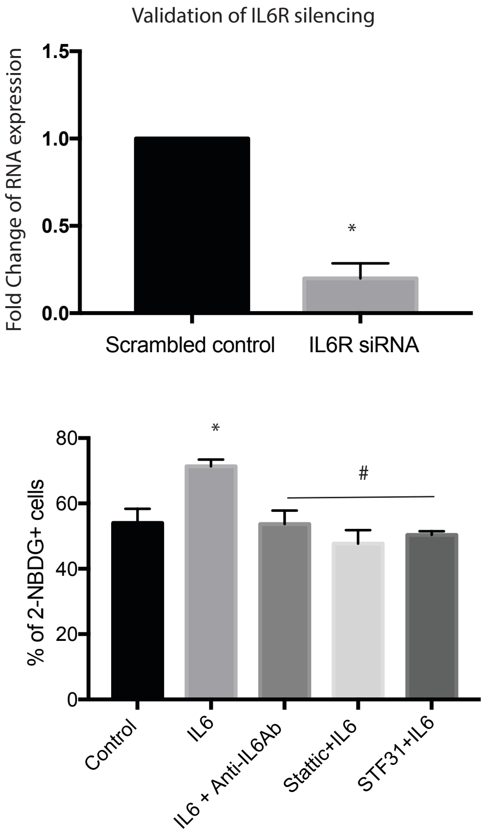
Validation of IL6R silencing
Fold Change of RNA expression
1.5
1.0
0.5
0.0
*
Scrambled control
IL6R siRNA
% of 2-NBDG+ cells
80
60
40
20
0
*
#
Control
IL6
IL6 + Anti-IL6Ab
Stattic+IL6
STF31+IL6

Annexin V Assay on Live Cells

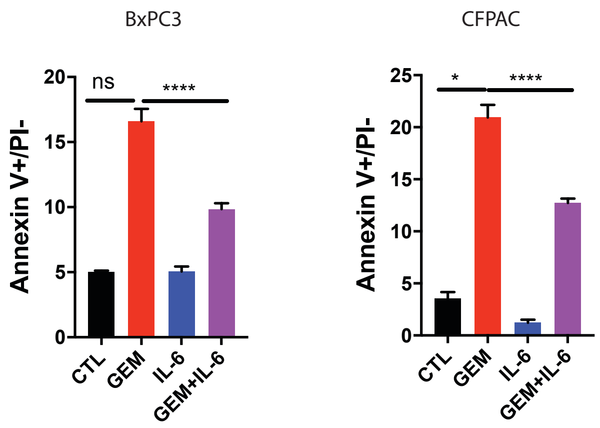


Supplementary Figure 1: Treatment of pancreatic cancer cells with IL6 increased their sensitivity to Gemcitabine as seen by decreased Annexin V+ cells in the apoptotic cell death assay.

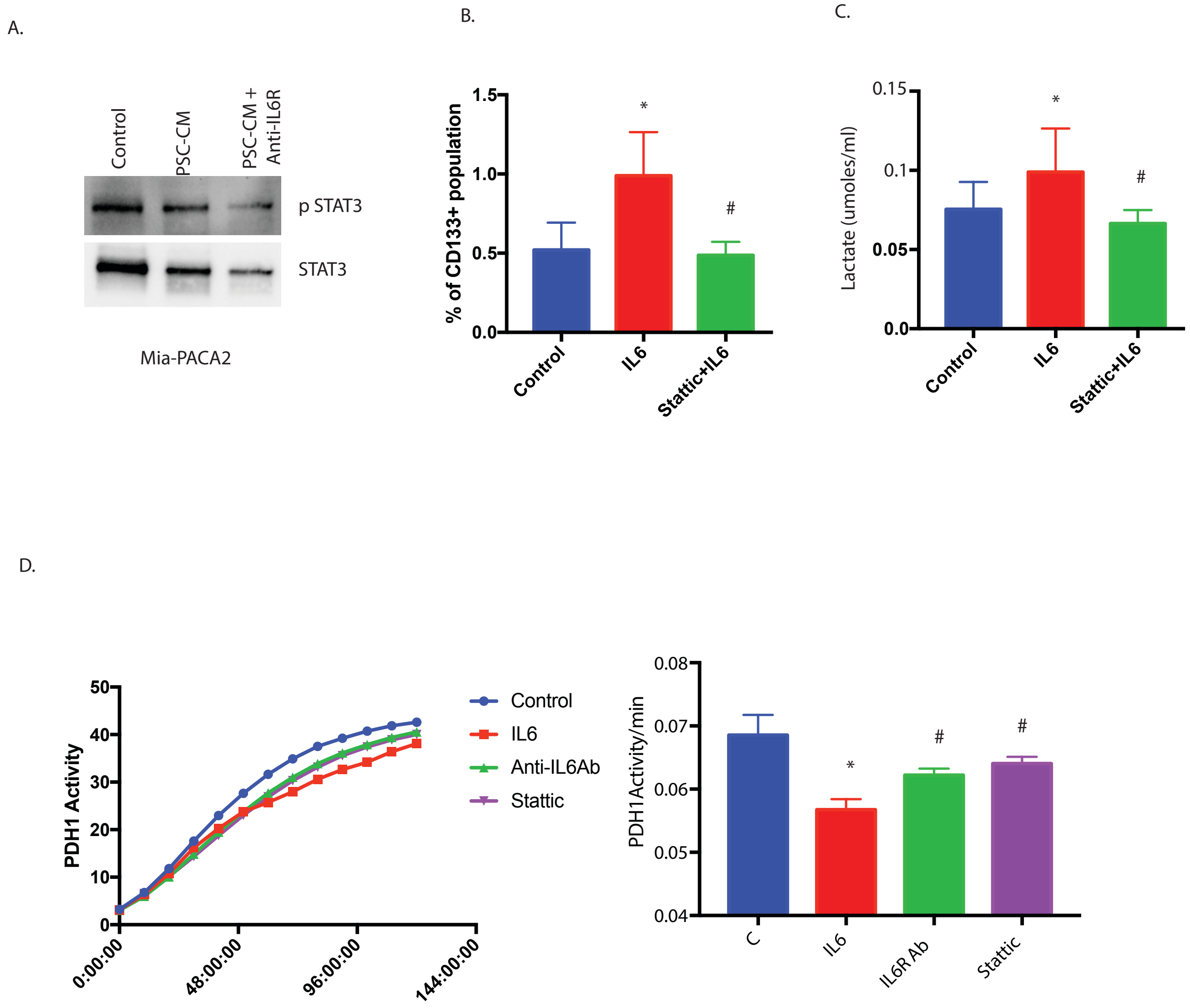

A.
Control
PSC-CM
PSC-CM + Anti-IL6R
p STAT3
STAT3
Mia-PACA2
B.
% of CD133+ population
1.5
1.0
0.5
0.0
*
#
Control
IL6
Stattic+IL6
C.
Lactate (umoles/ml)
0.15
0.1
0.05
0.0
*
#
Control
IL6
Stattic+IL6
D.
PDH1 Activity
50
40
30
20
10
0
0:00:00
48:00:00
96:00:00
144:00:00
Control
IL6
Anti-IL6Ab
Stattic
PDH1Activity/min
0.08
0.07
0.06
0.05
0.04
*
#
#
C
IL6
IL6R Ab
Stattic

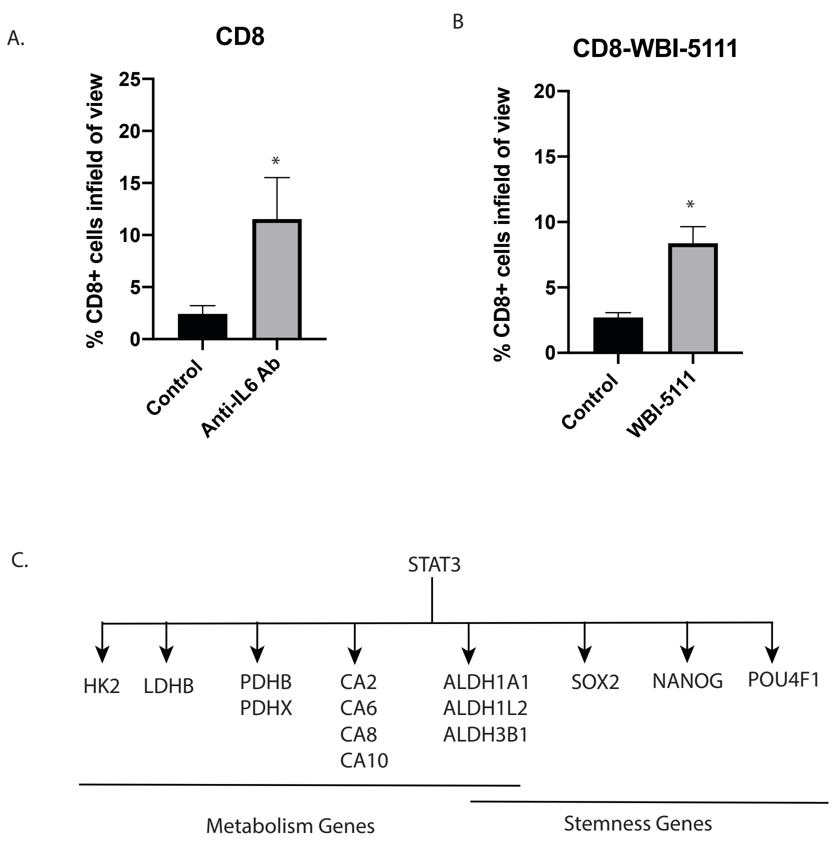


Analysis from CHEA Transcription Factor Targets
